# RnR-ExM: Robust Non-rigid Registration Challenge for Expansion Microscopy Volumes

**DOI:** 10.64898/2026.10.22.760871

**Authors:** Emma Besier, Ruihan Zhang, Yosuke Bando, Mansour Alawi, Yves Quéméner, Chaitanya Kapoor, Malte Hoffmann, Adrian Dalca, Adrià Casamitjana, Ignacio Arganda-Carreras, Greg Fleishman, Xi Jia, Jinming Duan, Tony C. W. Mok, Zi Li, Margaret E. Schroeder, Edward S. Boyden, Hanspeter Pfister, Donglai Wei

## Abstract

Expansion Microscopy (ExM) enables nanoscale imaging of biological samples while preserving structural and molecular details. The Robust Non-rigid Registration Challenge for ExM (RnR-ExM) was launched to benchmark 3D registration algorithms on real-world ExM datasets. The challenge released 24 pairs of high-resolution 3D volumes from zebrafish, mouse, and *C. elegans*, deliberately selected for their difficulty with standard alignment methods. To evaluate performance, we provided curated segmentation masks of nuclei and blood vessels as landmarks for half of the dataset. To encourage learning-based approaches, we released code for generating synthetic training data reflecting the deformation and appearance of the unlabeled pairs. By submission, 168 participants registered, with six teams submitting results by the deadline; the top three were invited to present at ISBI 2023. Accurate registration of ExM volumes is essential for downstream biological analyses such as multiplexed molecular mapping and cell-type-specific structural profiling. In this paper, we detail the methods and results of these top teams, highlighting current capabilities and future directions for 3D registration in ExM. All datasets and code are available at https://rnr-exm.grand-challenge.org/

## 1 Introduction

Expansion Microscopy (ExM) is a transformative imaging technique that enables nanoscale resolution using conventional fluorescence microscopes by physically expanding biological specimens embedded in swellable polymers [1]. When paired with multiplexed staining protocols, ExM facilitates the spatially resolved analysis of numerous molecular targets, ranging from nucleic acids to proteins, within intact 3D tissues [2]. However, the repeated staining and imaging rounds introduce significant non-linear deformations, often driven by gel movement or handling artifacts. Thus, accurate alignment of volumetric images across rounds becomes a critical yet unresolved challenge for downstream analysis, such as protein co-localization or cell-type-specific structure mapping.

Despite advances in medical image registration, existing benchmarks and competitions—such as OASIS [3], ANHIR [4], and Learn2Reg [5]—primarily target clinical imaging modalities like MRI, CT, and histology. These efforts do not address the unique challenges of ExM, which include nanoscale tissue deformations, large non-linear displacements, variable staining contrast, and limited annotation within high-resolution volumetric data. Additionally, there are currently no publicly available datasets for systematically evaluating registration performance under these ExM-specific conditions. Without accurate registration, ExM-based analyses risk losing nanoscale spatial fidelity across imaging rounds, compromising current efforts in synaptic protein mapping, high-resolution transcriptomics (*e*.*g*., ExM+MERFISH), and multi-round expansion workflows applied to brain and tumor tissues.

To address this gap, we introduce the *Robust Non-rigid Registration for Expansion Microscopy* (RnR-ExM) challenge, the first benchmarking initiative dedicated to evaluating registration algorithms on real-world ExM data with large deformations and sparse but biologically meaningful ground truth. The challenge was hosted at the 2023 IEEE International Symposium on Biomedical Imaging (ISBI), and features:

- A curated benchmark dataset of 24 3D volume pairs spanning three species (zebrafish, mouse, *C. elegans*),
- Sparse but biologically meaningful correspondence annotations (nuclei and blood vessels),
- A standardized evaluation pipeline with established registration metrics (*e*.*g*., Dice, deformation smoothness), and
- Open-source tools to generate synthetic training data mirroring real ExM deformations.

Following the live challenge, we conducted a post-hoc analysis to re-evaluate results at full resolution and extend the leaderboard with new metrics and insights that would not have been feasible during the initial timeline.

### Broader Impact

The RnR-ExM challenge was designed not only to benchmark registration algorithms, but also to expose key algorithmic and practical limitations in aligning ExM volumes—including failure cases in specific anatomical regions, trade-offs between runtime and accuracy, and difficulty generalizing across diverse tissue types. This paper presents the dataset and challenge infrastructure, highlights results from top-performing teams, and offers an in-depth analysis of their performance under ExM-specific constraints. Our aim is to provide a reusable benchmark and actionable insights to support the development of more scalable, robust 3D registration algorithms.

Importantly, the challenge addresses a growing need in biomedical imaging. As ExM becomes foundational to spatial proteomics, connectomics, and multiplexed imaging, reliable non-rigid registration is essential for maintaining spatial fidelity across imaging rounds. Misalignment can compromise molecular co-localization, distort cell-type-specific analyses, and obscure structural relationships. By enabling rigorous, reproducible comparison of methods under realistic ExM conditions, RnR-ExM supports the development of more accurate computational tools critical for downstream biological discovery in neuroscience, developmental biology, and beyond.

## 2 Challenge

### 2.1 Challenge Overview

Participants were tasked with aligning multiple pairs of 3D light microscopy volumes by estimating and submitting the corresponding deformation fields. Each deformation field represents a 3D displacement map that transforms the moving volume into alignment with the fixed reference volume by mapping corresponding spatial locations between them. The specific registration challenges associated with each species are summarized in Table 1.

**Table 1:**
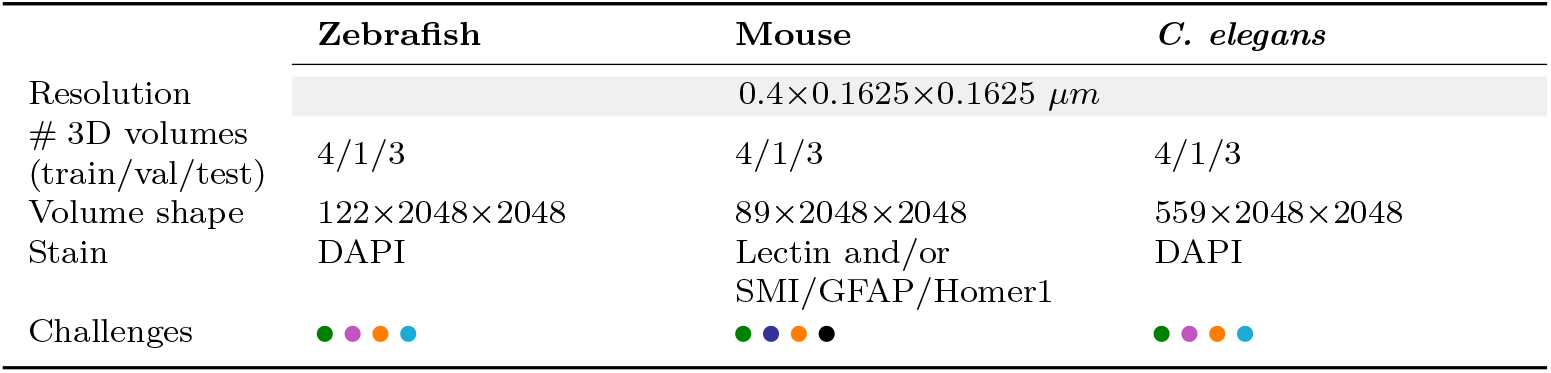
Overview of the RnR-ExM challenge, which shows a diverse set of ExM image registration difficulties: 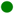 varying fields of view across image pairs, 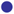 degraded image quality between rounds due to re-staining, 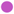 large deformation in the *z*-direction, 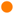 global linear deformations, 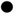 complex non-linear deformations, and 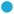 large image sizes.

**Table 2:** Best submission per team: *zebrafish*. Bolded values indicate the best score per metric.

| Team | DSC (↑) | DSC30 (↑) | HD95 (↓) | SDlogJ (↓) |
| --- | --- | --- | --- | --- |
| NLi10Me | 0.9721 | **0.9715** | 2.2056 | **0.0000** |
| Orange | **0.9730** | 0.9711 | **2.1911** | 0.0003 |
| Xi | 0.9707 | 0.9695 | 2.3028 | 0.0001 |

**Table 3:** Best submission per team: *mouse*. Bolded values indicate the best score per metric.

| Team | DSC (↑) | DSC30 (↑) | HD95 (↓) | SDlogJ (↓) |
| --- | --- | --- | --- | --- |
| NLi10Me | 0.8920 | 0.8383 | 10.7418 | **0.0000** |
| Orange | **0.9181** | **0.8684** | **7.8166** | 0.0464 |
| Xi | 0.8829 | 0.8576 | 13.1514 | **0.0000** |

**Table 4:**
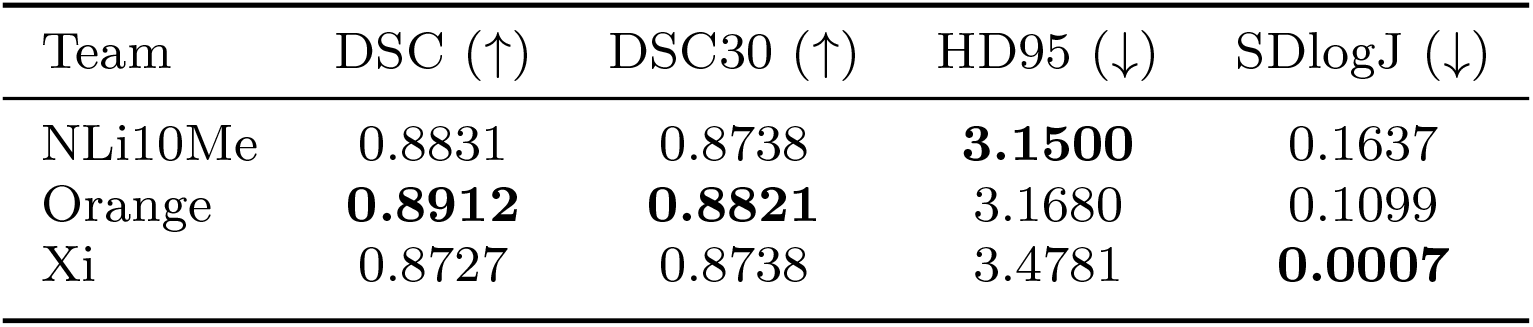
Best submission per team: *C. elegans*. Bolded values indicate the best score per metric.

### 2.2 Organization

#### Challenge Phases

The challenge comprised four phases:

- *Training phase (11/19/22 - 1/19/23)* Release of training set, code to generate synthetic dataset, baseline method and tutorial. The training set includes four volume pairs per species with ground truth annotations hosted at https://zenodo.org/records/7444100.
- *Validation phase (1/19/23 - 2/8/23)* Release of validation set and code for evaluation. The validation set includes one volume pair per species with ground truth annotations hosted at https://zenodo.org/records/8418961.
- *Test phase (2/8/23 - 2/15/23)* Release of test set images without ground truth annotations. The test set includes three volume pairs per species hosted at https://zenodo.org/records/8419002.
- *Evaluation phase (2/15/23 - 4/15/23)* After all data was released, participants were free to submit to the online leaderboard. Submission for ISBI 2023 closed on 4/15/23, but the online leaderboard remained open for submission past that date for continuous benchmarking.

#### Website

The challenge was hosted and available through Grand Challenge, “a platform for end-to-end development of machine learning solutions in biomedical imaging” [6]. Grand Challenge provides all of the functionalities for running a computing challenge: user management, access control, discussion forums, and support for creating leaderboards with custom evaluation algorithms. Grand Challenge agrees to host the challenge website and leaderboards for five years.

#### Post-Challenge Analysis

We undertook an extensive post-competition effort to enhance the reproducibility, depth, and analytical value of the results. The additional time between the live challenge and this manuscript has enabled a deeper and more generalizable analysis that would not have been possible during the original event timeline.

### 2.3 Benchmark Dataset

Our curated benchmark dataset contains 24 volumetric registration pairs from three commonly imaged species in neuroscience: zebrafish (brain), mouse (brain), and *C. elegans* (whole). Of these, twelve of the volume pairs have sparse manual labels for structure correspondence. For each species, the train, validation, test split is 4 : 1 : 3.

#### Data Acquisition

All data included in the competition dataset were collected using the data acquisition protocol detailed in [1]. We selected DAPI as the anchoring channel for the zebrafish and *C. elegans*, and Lectin or Lectin plus glial fibrilarly acidic protein (GFAP), neurofilament (SMI), and post-synaptic scaffold protein Homer1 as the anchoring channel for the mouse. We note that Lectin is the dominant (in terms of signal intensity and volume) reference stain in the combined mouse reference channel used in later imaging rounds. DAPI and Lectin are common stains across labs, are compatible with most experimental conditions and provide salient features. The size of each volume is *z ×* 2048 *×* 2048 voxels (*z ∈ {*89, 122, 559*}*) at a resolution of 0.4 *×* 0.1625 *×* 0.1625 *μ*m^3^.

#### Gold Standard Establishment

After choosing and compiling the raw image volumes (Fig. 1), we segmented the nuclei (zebrafish, *C. elegans*) and blood vessels (mouse) and found their structural matches across rounds. These structural matches were then used for accuracy evaluation as described in Sec. 2.4. Given the size of our dataset, the annotations are sparse (*e*.*g*., not all structures are annotated) and are only available for half of all volumes (the validation and test sets). An overview of the segmentation method used for each species is outlined below. Examples of sparse segmentation masks with matched landmarks are shown in Figure 2.

**Fig. 1:**
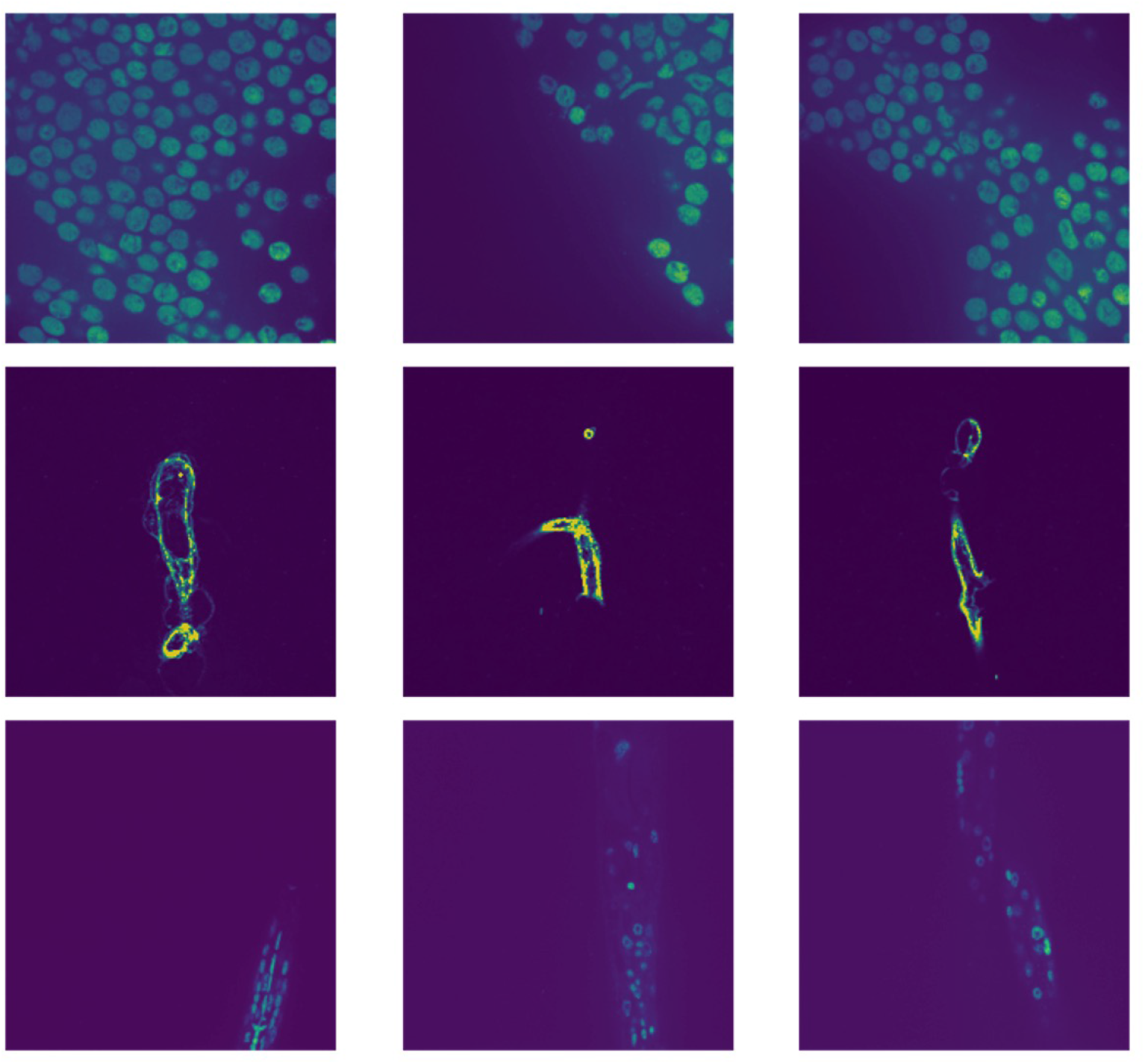
Example *z*-slices from each of the three species. zebrafish (**top**), mouse (**middle**), and *C. elegans* (**bottom**). The circular reference structures visible in the zebrafish and *C. elegans* slices are nuclei, and the reference structures visible in the mouse slices are blood vessels.

**Fig. 2:**
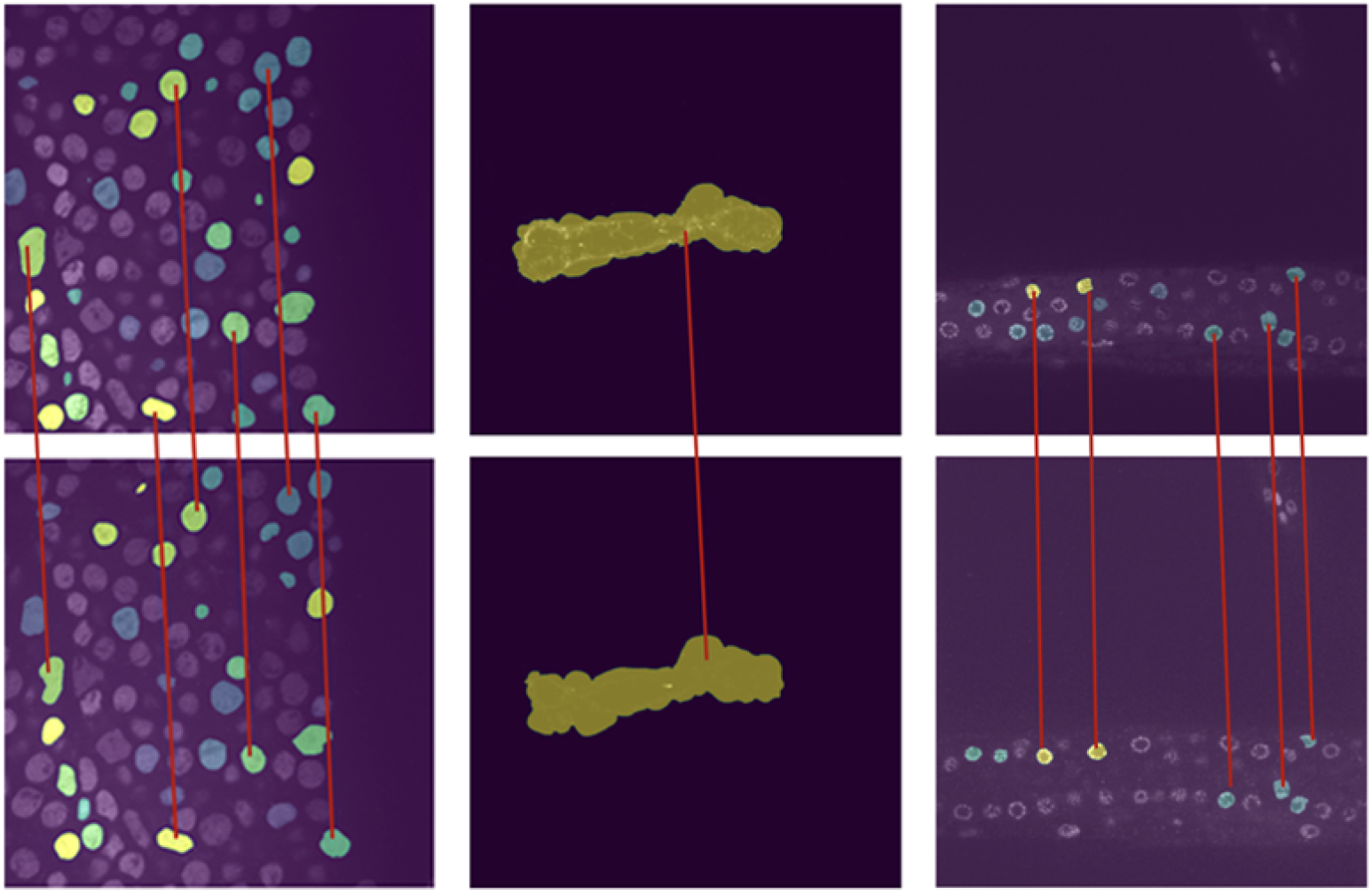
Examples of sparse segmentation masks overlaid on original volume slices. zebrafish (**top**), mouse (**middle**), and *C. elegans* (**bottom**). The fixed volume slice is shown on the left, and the corresponding slice from the moving volume is on the right. Matched structures are labeled across both volumes.

- *Zebrafish:* To segment zebrafish nuclei, we employed a fine-tuned version of Cellpose [7], a generalist segmentation model capable of transferring annotations across cellular imaging domains. The resulting masks were manually inspected and refined using the VAST annotation tool [8], followed by median filtering to smooth label boundaries. Corresponding nuclei across imaging rounds were assigned consistent labels, while all unmatched regions were set to zero. Manual matches were independently reviewed by three annotators, and any instances of disagreement were excluded from the final annotations. We manually verified 43, 29, and 61 pairs of matched nuclei for the train, validation, and test splits, respectively.
- *Mouse:* Each mouse volume contained a single prominent feature: a blood vessel. To segment this structure, we first performed foreground-background separation using manual thresholding. We then applied the OpenCV [9] findContours function to isolate and fill the primary landmark while removing background noise. The resulting masks were manual refined in the VAST to ensure accurate structure delineation across volume pairs.We manually verified one pair of matched nuclei for the train, validation, and test splits, respectively.
- *C. elegans:* Multiple automated segmentation methods were evaluated, but none proved effective in reliably identifying and separating nuclei in three dimensions. A key challenge was the presence of nuclei with poorly defined boundaries, which hindered algorithmic detection. As a result, segmentation and matching were performed manually by multiple annotators in the VAST.We manually verified 42, 72, and 66 pairs of matched nuclei for the train, validation, and test splits, respectively.

To facilitate the development of learning-based methods, we released open-source code for generating synthetic training data. The pipeline simulates realistic ExM deformations by randomly sampling transformation parameters that emulate either unintentional drift of the microscope objective or intrinsic sample deformation. Each synthetic transformation consists of a large global affine component followed by a finer piecewise affine deformation. Deformation fields are constructed by sampling random motion vectors on a regular grid and applying Gaussian smoothing. This approach enables the generation of an unlimited number of synthetic volume pairs that closely approximate the deformation characteristics and visual appearance of the unlabeled challenge data.

### 2.4 Evaluation

#### Evaluation metrics

Following recent registration challenges [5], we evaluated the accuracy of each submitted algorithm (in the form of deformation fields) based on segmentation labels. Specifically, the segmentation labels of each moving volume were transformed by the deformation field, and the evaluation was based on how well the deformed segmentation labels aligned with those of the corresponding fixed volume. The following metrics were used for assessing the quality of the alignment.

- *DSC*: The Dice similarity coefficient [10] measures the overlap between two sets of segmentation labels, calculated as 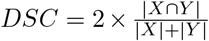, where *X* and *Y* are the segmentation labels of the fixed and moving volumes.
- *DSC30*: The 30^th^ percentile of the DSC values taken over all volume pairs assesses algorithm robustness.
- *HD95*: The Hausdorff distance measures the maximum distance between the 3D surfaces of corresponding segmentation labels, defined as HD(*X, Y*) = {max max_*x∈X*_ min_*y∈Y*_ *d*(*x, y*)}, max_*y∈Y*_ min_*x∈X*_ *d*(*x, y*), where *d*(*x, y*) is the Euclidean distance between points *x* and *y* in segmentation labels *X* and *Y*. HD95 takes the 95^th^ percentile instead of the maximum of these distances to reduce the impact of outliers as a robust measure of the maximum mismatch between surfaces.
- *SDlogJ*: The standard deviation of the logarithm of the Jacobian determinant of each deformation field, calculated as 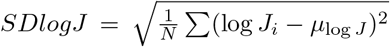, is used to assess deformation smoothness, where *J*_*i*_ is the Jacobian determinant at each voxel, *μ*_log *J*_ is the mean log Jacobian across the volume, and *N* is the total number of voxels. Non-smooth deformation is not realistic, and this metric penalizes such submissions.

#### Ranking Mechanism

To determine the leaderboard standings, we combined all evaluation metrics into a unified ranking score following the methodology in [11]. For each submission, we calculated its *significance rank*, defined as the number of competing algorithms that performed statistically significantly worse under a given metric. A higher significance rank thus indicates superior performance. These ranks were then normalized and averaged across all volume pairs and evaluation metrics to produce a single aggregate score per algorithm. For each task, we maintained two leaderboards: a full leaderboard displaying all submissions, and a reduced leaderboard showing only the top submission from each team.

#### Self-reported runtime

This competition did not require the release of code, however participants were asked to submit a speed assessment that lists the runtime of each algorithm as well as information about the host environment (machine type, the number of CPU cores or GPUs used, and the clock frequency each processing unit supports). Because we had no way of enforcing the truthful reporting of speed, speed was not used to determine position on the leaderboard.

### 2.5 Submitted Methods

To provide a reference point for evaluation, we released a baseline registration method using an out-of-the-box rigid transformation model implemented in SimpleITK [15]. The baseline serves both as a minimal benchmark and a framework for reproducibility. Configuration included standard parameters—rigid transformation type, mutual information similarity metric, default optimizer and interpolator—with resolution set to match the physical voxel spacing of each image volume.

The challenge attracted significant interest, with 137 participants registering on the Grand Challenge platform. Of these, six teams submitted valid results, yielding a total of 96 submissions. Final leaderboard rankings were computed after re-evaluation at full resolution. The top three teams were invited to present at the ISBI 2023 challenge workshop and contribute detailed method descriptions for this manuscript. Their approaches represent a diverse mix of optimization-based and learning-based pipelines, and are analyzed in detail in the following sections.

## 3 Results

### 3.1 Top-3 Participant Methods

In this section we discuss the methods of the three top-performing teams: *NLi10Me* (first place), *Orange* (second place), *Xi* (third place). The top three teams are featured because there was a significant dropoff in performance between third and fourth place submissions. The accompanying code and publications for each of the methods are listed in Table 5.

**Table 5:** Summary of top-3 participant methods.

| Participant | Method | Type | Code | Publication |
| --- | --- | --- | --- | --- |
| NLi10Me | BigStream | Non-learning | Link | [12] |
| Orange | Similarity Pyramids | Non-learning | – | [13] |
| Xi | LKU-Net-Affine | Learning | Link | [14] |

- *NLi10Me* used BigStream [16], a registration toolkit for large-scale 3D image alignment. To handle volumes exceeding memory or thread limits, BigStream partitions data into overlapping blocks, processes them in parallel using Dask [17], and stitches the results into a smooth global transform. The pipeline combined affine and optional deformable models implemented via SimpleITK [15]. All alignments used gradient descent on mutual information with linear interpolation, and volumes were downsampled to reduce runtime. Deformable components were modeled using cubic *b*-splines with multiscale control point optimization.
- *Orange* employed a two-step registration pipeline comprising affine alignment followed by optional non-rigid optimization. The affine step initialized with an identity matrix and iteratively updated the transformation using image pyramids and normalized cross-correlation (NCC) [18]. For volumes exhibiting non-linear deformations, a non-rigid refinement step was applied using similarity pyramids [19], with a diffusion regularizer added to promote smooth and plausible deformations.
- *Xi* used LKU-Net-Affine, a deep learning model extending the LKU-Net encoder [14] with an additional affine prediction head comprising four modules of fully connected layers. The model was trained using online 3D data augmentation to address limited annotated data, and optimized using locally normalized cross-correlation (LNCC) as the similarity metric. Final predictions were further fine-tuned individually on each test volume pair to improve alignment precision.

#### Method Novelty

While the core architectures used by the top teams were adapted from existing frameworks, each method exhibited notable engineering innovations that made them suitable for the unique constraints of ExM data. These innovations spanned system-level scalability, resource-aware deployment, and data-efficient learning—demonstrating that practical novelty often lies in bridging algorithm design with real-world usability.

*NLi10Me*’s BigStream addressed the challenge of terabyte-scale volume alignment by automating blockwise decomposition, parallel registration, and smooth stitching. This pipeline supports both affine and deformable models and enables alignment of datasets that exceed GPU or RAM capacity—critical for large-tissue ExM imaging.

*Orange* showed that a classical optimization pipeline, when carefully tuned, can remain competitive without deep learning. Their method leveraged multi-scale affine alignment, normalized cross-correlation, and diffusion regularization, all implemented with low computational overhead—making it ideal for deployment in non-GPU environments.

*Xi* extended a UNet encoder with an affine prediction head and used online synthetic augmentation and test-time fine-tuning to overcome annotation scarcity. Though composed of standard modules, this architecture adapted effectively to the visual heterogeneity of ExM volumes and achieved strong results under limited supervision.

### 3.2 Method Comparisons

Dice scores were comparable across all three methods, with the greatest variability observed in the mouse data (ranging from 0.892 to 0.918). The largest differences between teams emerged in the HD95 and SDlogJ metrics. Table **??** summarizes each team’s best scores, averaged over the three test pairs.

#### Performance on Each Volume

Figure 3 shows performance broken down by individual test pair. As outlined in Sec. 2.4, final rankings were computed using a significance-based scoring system that considers all evaluation metrics collectively. This explains why a team may not achieve the highest overall rank despite outperforming others on specific metrics.

**Fig. 3:**
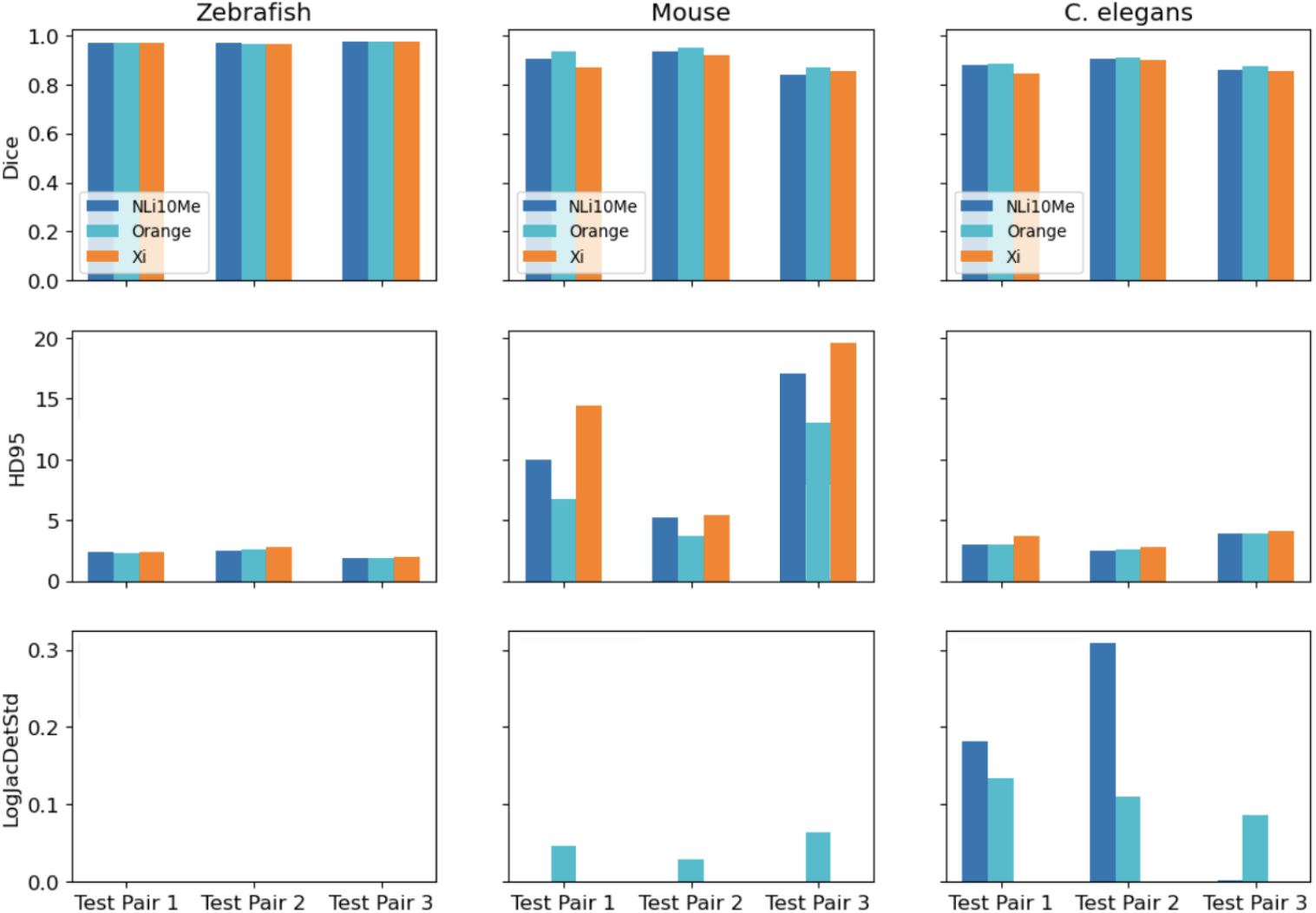
Test performance of top-3 participant methods.

#### Performance on Each *z*-Slice

Next, we investigated whether there was a specific volume region that was consistently difficult for participants to align. This involved plotting the error (DSC) across all *z*-slices and across nuclei. No consistent error patterns were found when plotting across *z*-slices (Figure 4), eliminating conclusions suggesting a *z*-dependence in registration quality or difficulty.

**Fig. 4:**
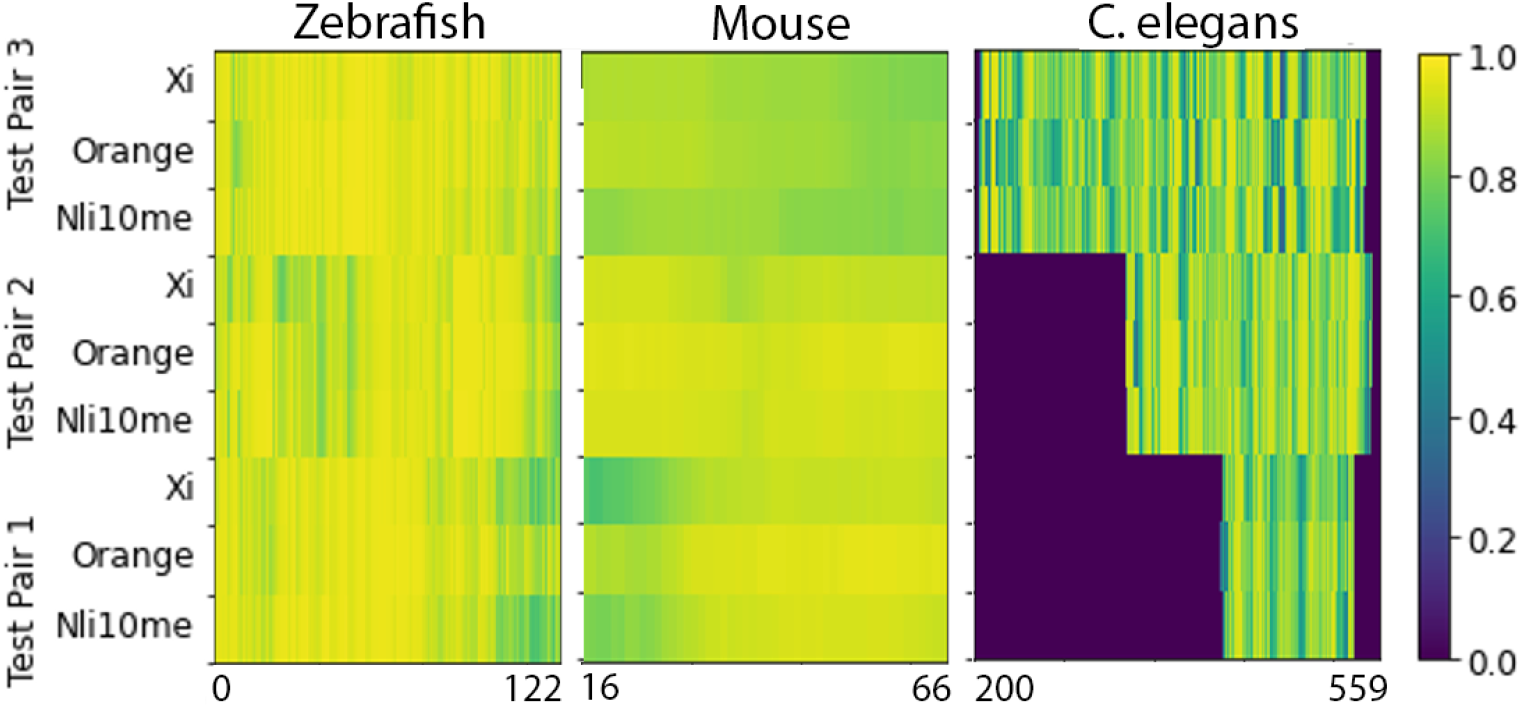
DSC on each test *z*-slice. *C. elegans* data contains empty slices in both ends of the volumes with zero DSC.

#### Performance on Selected Nuclei

We further analyzed DSC values across *z*-slices for a set of randomly selected nuclei in the zebrafish test data (Figure 5). Across all participants, the highest alignment errors consistently appeared at the start and end of each nucleus. This may stem from the lower resolution in the *z*-axis compared to the *x*- and *y*-axes, and the smaller cross-sectional area of nuclei at their boundaries. In such regions, even small misalignments can cause a disproportionately large drop in DSC, while similar displacements in central slices have less impact due to larger structure size.

**Fig. 5:**
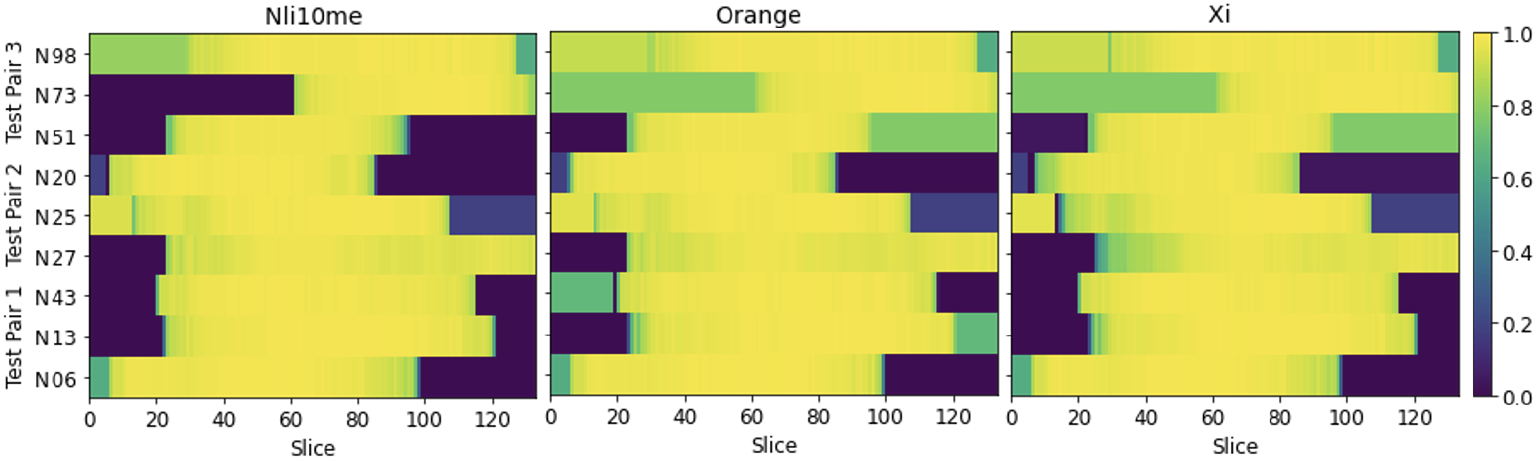
DSC on selected nuclei. The highest alignment errors consistently occur at the start and end of the nuclei, as indicated by lower DSC values at the beginning and end of each row. This pattern is observed across all participants (**left**: NLi10Me, **center**: Orange, **right**: Xi). Zebrafish was chosen for this analysis due to the use of a fine-tuned deep learning model for segmentation, minimizing annotation variability.

#### Failure Cases

We used three different plots to further assess the quality of submissions and to identify the remaining bottlenecks in ExM registration. First, we examined slices with high alignment error (low DSC) by overlaying the fixed and transformed moving segmentation masks (Figure 6). The mouse and *C. elegans* volumes exhibited the most pronounced visible errors in these worst-case slices, with discrepancies substantial enough to affect downstream analyses. Notably, in the *C. elegans* data, we observed complete mismatches between nuclei in the fixed and moving masks. Further inspection revealed that these failures often occurred in regions with densely packed or overlapping nuclei, weak edge contrast, and at sites of nucleus splitting or merging across slices. In the mouse data, errors were most pronounced near vessel bifurcations and regions with partial volume effects, where the vessel boundary becomes ambiguous due to reduced staining intensity or sharp curvature. These findings highlight that anatomical complexities, such as closely apposed structures and branching geometries, remain challenging for current registration methods, indicating areas for targeted algorithmic improvement in future ExM registration pipelines.

**Fig. 6:**
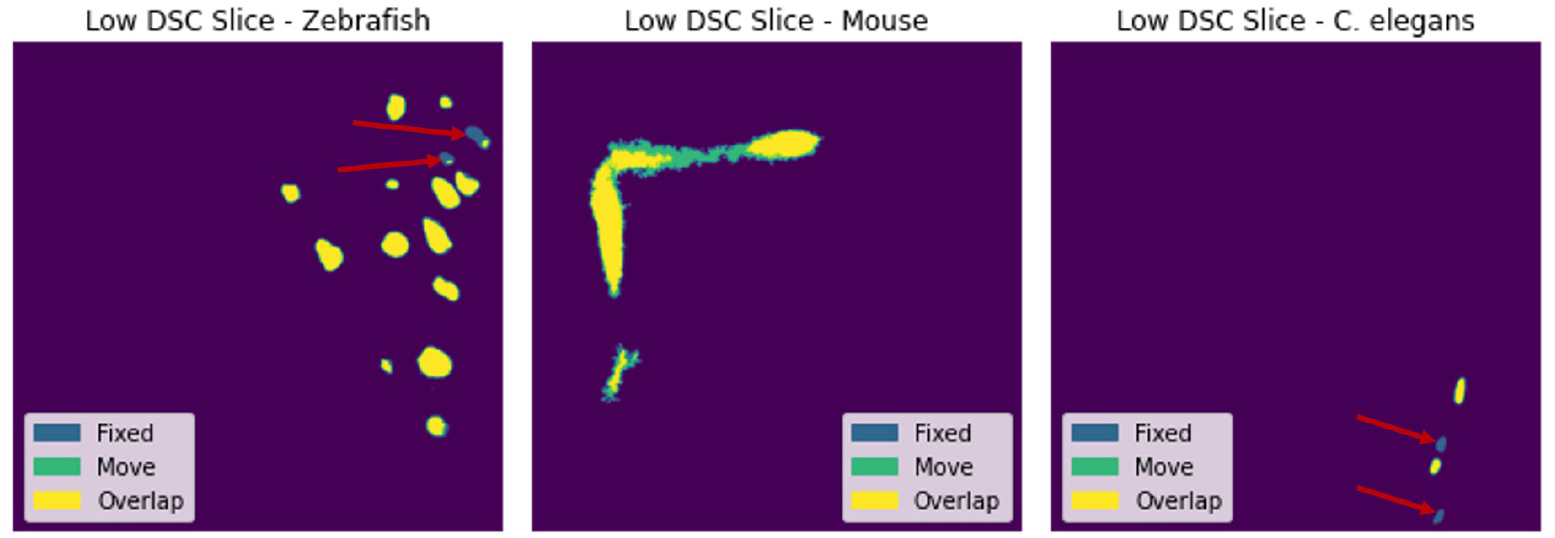
Failure cases. Low-DSC-scoring fixed segmentation masks overlaid with transformed moving segmentation masks: zebrafish (left, 0.77), mouse (center, 0.86) and *C. elegans* (right, 0.46). Arrows indicates major mismatches.

#### Reported Runtimes

Participant-reported runtimes are summarized in Table 6 with specified machine information. Notably, two teams—*NLi10Me* and *Xi* —included I/O and downsampling time in their measurements, while *Orange* did not. *NLi10Me*’s most accurate *C. elegans* submission required 9,200 seconds, due to the use of small processing chunks aimed at maximizing alignment accuracy. While larger chunks would have reduced runtime, this would have compromised performance, particularly in terms of DSC. Similarly, *Xi*, which employed a learning-based method, was constrained to a maximum image resolution of *z ×* 256 *×* 256 for inference. The team noted that using higher-resolution inputs would have improved accuracy, but exceeded runtime budgets.These findings reveal a trade-off between computational efficiency and alignment precision in ExM registration.

**Table 6:**
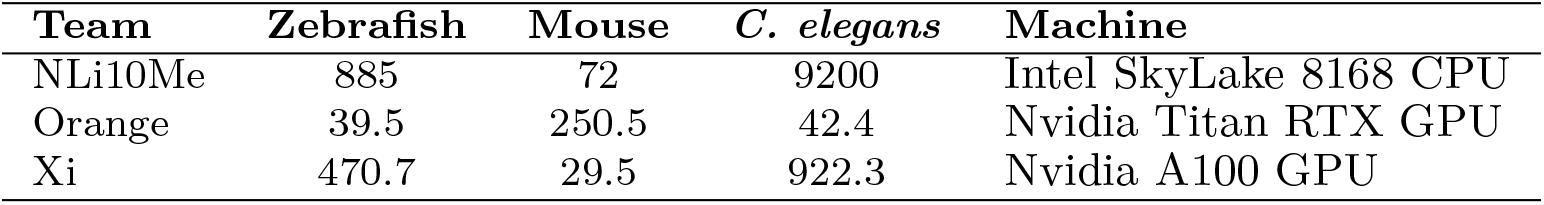
Reported runtimes in seconds.

#### Model generalization

One benefit of learning-based methods is that once trained, they can quickly predict a deformation field between a pair of volumes. To be effective, machine learning methods require model generalization. *Xi*, the team that used a learning-based approach, reported that the model was not able to generalize beyond the training data, even though the volumes were gathered from the same experiment. Thus, *Xi* fine-tuned (manually tweaked) the model hyper-parameters for each pair in the test set. This led to runtimes that were occasionally greater than non-learning based approaches. It is possible that training with more data would resolve this issue. Synthetic data generation was incorporated into the pipeline, but it did not aid in model generalization.

#### Method accessibility

An important consideration is algorithm accessibility. ExM registration is both computationally-expensive and difficult to troubleshoot. In bio-focused labs, it is often biologists familiar with coding that handle the computational work, not trained software engineers. Registration pipelines should be accessible to an audience with limited coding knowledge. Notably, *NLi10Me*’s method (BigStream) is built on and uses syntax akin to SimpleITK [15], a user-friendly registration package with a number of beginner tutorials online. It is not clear how accessible the other discussed methods are. *Xi* ‘s method requires model training and fine-tuning, which may not be intuitive to bio-focused users. *Orange*’s code is not publicly available.

## 4 Discussion

### 4.1 Algorithm Design Strategies

Despite architectural differences, all three methods shared a common high-level structure tailored to the demands of ExM registration. Each employed a global-to-local alignment pipeline to handle large deformations while maintaining anatomical accuracy. *NLi10Me* and *Orange* explicitly performed multi-resolution affine initialization, while *Xi* implemented this stage internally via its learned affine header. This coarse-to-fine design helped mitigate large initial misalignments and improved convergence stability.

Regularization strategies varied significantly and reflected different philosophies of deformation control. *Orange* penalized high-frequency motion using a diffusion term, effective in preserving smooth transitions across tissue but less flexible in locally deformed regions. *NLi10Me* maintained consistency by enforcing overlap across block-wise sub-volumes, which provided robustness but introduced edge-sensitive stitching artifacts. *Xi* ‘s approach was entirely implicit, relying on training data, network structure, and augmentations to discourage irregular deformations.

Together, these strategies illustrate that successful ExM registration requires balancing global alignment, local flexibility, and deformation plausibility—each team operationalized this balance differently, leading to variations in both strengths and failure modes across datasets.

### 4.2 Algorithm Failure Modes

Across all submissions, consistent error patterns emerged in regions with ambiguous or degraded visual cues—particularly low-contrast boundaries, fragmented structures, or anisotropic resolution. For example, bifurcating vessels in the mouse and edge-slice nuclei in *C. elegans* often led to visible misalignments. These errors disproportionately affected Dice scores due to the sparsity and shape variability of the annotated regions, especially where cross-sectional area was small or rapidly changing along the *z*-axis.

A closer analysis reveals that these failure cases were closely tied to the modeling assumptions and design trade-offs of each approach. Learning-based methods (e.g., *Xi*) relied on data-driven priors learned from synthetic examples, but these did not capture the full heterogeneity of real ExM image appearance. As a result, the model was prone to overfitting or misgeneralization in unseen contexts, such as faint or fragmented structures. Inference was further constrained by limited memory, forcing downsampling that blurred fine structures critical for accurate alignment.

Optimization-based methods (*NLi10Me, Orange*) were less affected by data distribution shifts, but their reliance on global similarity metrics made them vulnerable in localized regions with weak or inconsistent signal. For example, normalized crosscorrelation (NCC) and mutual information performed well in well-contrasted regions but failed to guide alignment accurately in areas with low signal-to-noise or tissue discontinuities. Additionally, block-based registration strategies introduced stitching artifacts when chunk boundaries crossed through complex anatomical regions, such as bifurcations.

### 4.3 Dataset and Evaluation Limitations

Determining the proper amount of data to release was a significant challenge. Releasing too little could hinder the development of learning-based methods, while releasing too much risked overwhelming teams without access to substantial computational resources. Even with our conservative data release, some participants required assistance with resource-intensive tasks, *e*.*g*.up-sampling their predicted deformation fields. Future iterations of the challenge should explore ways to increase dataset availability without sacrificing accessibility.

Evaluation posed another constraint, particularly due to the RAM limitations imposed by the Grand Challenge platform. To accommodate these restrictions, we downsampled submitted deformation fields in the containerized evaluation pipeline, which led us to exclude the SDlogJ metric from the live leaderboard—its downsampled version did not accurately reflect full-resolution behavior. After the competition concluded, all submissions were re-evaluated at full resolution on a local MIT server, incorporating SDlogJ into the final rankings. This post-hoc evaluation required significant manual effort and nearly a week of compute time. Alternative, more scalable evaluation strategies should be considered for future editions.

### 4.4 Future Directions

While the RnR-ExM challenge establishes a foundational benchmark for 3D registration in Expansion Microscopy (ExM), there remain numerous avenues for advancing both algorithmic performance and practical utility.

#### Handling Larger and More Complex Deformations

Future benchmarks should include more complex registration scenarios, such as aligning pre- and post-expansion images or performing cross-modality registration between ExM and electron microscopy. Such tasks demand methods that can model highly non-linear, spatially heterogeneous tissue deformation patterns that go beyond simple affine or smooth warping fields.

#### Generalization in Learning-Based Methods

In the RnR-ExM challenge, deep models required fine-tuning for each test volume, which limits their scalability. A promising direction is to develop models that leverage domain adaptation, semi-supervised learning, or self-supervised objectives to improve generalization without re-training. In parallel, synthetic data augmentation strategies could be refined to better reflect the full spectrum of biological and imaging variations encountered in ExM, enhancing their utility for model pretraining.

#### *z*-axis Resolution Adaptivity

Future methods should explicitly model this anisotropy, for example, through architectures that adapt to *z*-resolution or incorporate slice-aware regularization that balances smoothness with the preservation of biologically relevant discontinuities. Addressing this challenge is critical for accurately aligning extended or sparsely segmented structures such as elongated nuclei or blood vessels.

#### Runtime and Memory Efficiency

Current state-of-the-art methods require hours of processing time per volume, often relying on large-scale parallelism or downsampling. Next-generation models should explore scalable architectures that preserve accuracy while reducing runtime and memory overhead, possibly by leveraging hierarchical or attention-based strategies. Moreover, enabling users to control the trade-off between accuracy and speed at inference time would make these tools more adaptable to diverse experimental constraints.

#### Benchmark and Metric Expansion

Incorporating biologically motivated metrics, such as co-localization error, would provide a more meaningful assessment of registration quality in real-world applications. Including both densely annotated synthetic volumes and sparsely labeled real data could help measure performance across different annotation regimes. Furthermore, integrating uncertainty quantification into registration outputs would allow users to identify regions of potential misalignment and improve interpretability.

#### Accessibility and Deployment

To ensure adoption by biology-focused labs, methods should be packaged into user-friendly, low-code pipelines that can be integrated into existing image analysis platforms. These tools should be hardware-agnostic, supporting both CPU-only and GPU-based environments, and ideally offer interactive interfaces for refining alignments using user input such as landmarks or structural prompts. Enhancing usability in this way will be essential for bringing robust non-rigid registration tools from research settings into routine biological workflows.

## 5 Conclusion

The Robust Non-rigid Registration for Expansion Microscopy (RnR-ExM) challenge represents the first dedicated effort to benchmark 3D registration algorithms in the context of Expansion Microscopy—a setting marked by large, biologically induced non-linear deformations, sparse annotations, and high-resolution volumetric data. By assembling a curated dataset of 24 volume pairs across three species and designing a multi-phase, public evaluation framework, this challenge aims to catalyze algorithm development, facilitate comparative assessment, and set realistic expectations for registration performance under ExM-specific constraints.

Our analysis of the top-performing submissions highlights that both optimization-based and learning-based approaches can achieve comparable alignment accuracy when carefully tuned. However, persistent challenges remain: aligning small or sparsely segmented structures, handling anisotropic resolution (especially in the *z*-direction), balancing computational runtime with deformation fidelity, and achieving generalization without per-sample fine-tuning.

Beyond rankings, the RnR-ExM challenge establishes a reproducible and extensible benchmark infrastructure to support continued research in 3D registration for biological microscopy. It underscores the importance of method accessibility, inference efficiency, and biological validity, key factors that influence real-world adoption in biomedical labs. As multiplexed ExM protocols continue to scale in resolution and throughput, robust and efficient registration will be essential for downstream tasks such as spatial transcriptomics integration, organelle tracking, and tissue atlasing.

We envision RnR-ExM as a living benchmark that evolves with the field. Future extensions may include more challenging deformation regimes (*e*.*g*., pre-vs. post-expansion registration), richer annotations, and community-driven method contributions. In doing so, RnR-ExM provides a critical foundation for advancing robust, scalable, and biologically informed image registration in the era of high-content 3D microscopy.

## Acknowledgements

I.A-C. acknowledges grants PID2024-157485NB-I00, RED2024-153844-T and AIA2025-164165-C42 funded by MICIU/AEI/10.13039/501100011033 and by FEDER, and the financial support from Gobierno Vasco-Eusko Jaurlaritza (IT2001-26). M.E.S. acknowledges NSF GRFP #1745302, the MathWorks Science Fellowship at MIT, and the Collamore-Rogers Fellowship at MIT. E.S.B. acknowledges NIH 1R01EB024261, Lisa Yang, John Doerr, NIH 1R01MH123403, NIH 1R01MH123977, and Schmidt Futures. H.P. acknowledges NSF IIS-2124179 and NIH 5U54CA225088-03. D.W. acknowledges NSF IIS-2239688.

## Notes

https://rnr-exm.grand-challenge.org/

https://zenodo.org/records/7444100

https://zenodo.org/records/8418961

https://zenodo.org/records/8419002

### Competing Interest Statement

The authors have declared no competing interest.

